# Complexin-1 regulated transition in the assembly of single neuronal SNARE complex

**DOI:** 10.1101/2021.06.07.447331

**Authors:** Tongrui Hao, Nan Feng, Fan Gong, Jiaquan Liu, Lu Ma, Yu-Xuan Ren

**Author notes:** **Correspondence should be addressed to** (T. H.); (J. L.); (L. M.); (Y. R.).

## Abstract

Neurotransmitter release is mediated by the synaptic vesicle exocytosis. Important proteins in this process have been identified including the molecular machine Synaptic-soluble N-ethylmaleimide-sensitive factor attachment receptor (SNARE) proteins, and other regulators. Complexin (Cpx) is one of the vital regulators in this process. The functions of Cpx are proposed to maintain a proper primed vesicle pool by preventing its premature depletion, which facilitates the vesicle fusion in the presence of Ca^2+^. However, the molecular mechanism remains unclear. Using dual-trap optical tweezers, we detected the interaction of complexin-1 (CpxI) with SNARE. We found that the CpxI stabilizes partially folded SNARE complexes by competing with C-terminal of Vamp protein and interacting with the C-terminal of t-SNARE complex.

## INTRODUCTION

The neurotransmitter release and intercellular communication requires the membrane fusion mediated by synaptic-soluble N-ethylmaleimide-sensitive factor attachment receptor (SNARE) proteins. The membrane fusion usually completes within a millisecond upon trigger.^1^ During the process, many regulatory proteins, such as N-ethylmaleimide sensitive factor (NSF), soluble NSF adaptor proteins (SNAPs), Munc18-1, Munc13s, complexin (Cpx) and synaptotagmin-1, are involved to regulate the zippering of SNARE complex and guarantee that the membrane fusion occurs at the precise time with high efficiency in the cell^2, 3^.

The functions of some important regulators have not been well understood well, such as Cpx, a small cytosolic α-helical protein. The Cpx is abundant in the presynaptic terminal and was first identified due to its ability to bind to SNARE complexes^4^. Cpx might have two functions: act as a clamp to inhibit the release to maintain a proper primed vesicle pool^5^, and facilitate Ca^2+^-triggered synchronous neurotransmitter release. liposome assay^6^ and Hela cells that ectopically express “flipped” SNAREs on their cell surface showed that Cpx inhibits the SNARE-driven fusion machinery providing direct evidence for a negative modulatory role in exocytosis. Expression of either complexin-I or II markedly suppresses acetylcholine release from PC12 cells^7, 8^, and strongly impairs HGH secretion from insulin secreting cell lines^8^. These studies suggest Cpx might act as a clamp to inhibit the release to maintain a proper primed vesicle pool. What’s more, knock-out and knockdown studies of Cpx have shown a prominent reduction of evoked release, likely pointing to a direct facilitator role of Cpx in synchronous neurotransmitter release^9, 10^. While compromised evoked release may be due to depletion of primed vesicles by premature spontaneous fusion^11, 12^, this explanation isn’t in general applicable for all types of preparations. Single-vesicle content mixing and liposome fusion assays have provided convincing evidence for an enhanced Ca^2+^-control of vesicle fusion in the presence of Cpx^13, 14^ Thus, phenotypic cues from the vast majority of model systems as well as in vitro analyses indicate a fusion promoting action of Cpx that either complements concurrent Cpx-mediated “clamping” of spontaneous fusion or even represents its chief function depending on the particular model system^15^.

The molecular mechanism by which Cpx regulates SNARE assembly has long been controversial. This could be triggered by its multiple domains’ interaction with the SNARE complex. Cpx consists of four domains, N-terminal domain (NTD), accessory helix (AH), center helix (CH) and C-terminal domain (CTD). The crystal structure of Cpx/SNARE complex and mutagenesis experiments have revealed that the CH is directly associated with SNAREs, which is essential for Cpx function^16, 17^ Furthermore, the NTD promote the fusion process, while the accessory helix may enhance the binding between the central helix and SNARE, and inhibit the fully assembly of SNARE^18^. Lastly, the CTD is thought to position Cpx to synaptic vesicles in a curvature sensitive manner because this region contains an amphipathic helix that binds to phospholipids^19, 20^, thereby concentrating other inhibitory domains of Cpx (e.g., accessory helix) at the fusion site^20–22^; and have also been suspected to contribute to the stabilization of the central helix^23^. It has been shown to clamp spontaneous fusion in neurons^24–26^ and to hinder premature secretion in neuroendocrine cells^27^ While this form of proximity accelerated inhibition appears attractive to catalyze a reliable blockade of SNARE assembly, recent structure function analyses with C. elegans Cpx revealed that membrane binding is important but not sufficient for Cpx inhibitory effects^28, 29^. The interaction among multiple domains of Cpx is still unclear, and may help uncover the complex functions of Cpx.

Although the functions of Cpx have been extensively investigated, the question about how Cpx interacts with SNARE in functional state is still unsolved. In contradistinction, magnetic tweezers study has revealed that the unzipping of SNARE complexes occurs at higher force levels by ~2 pN on average by adding CpxI to the mechanical unzipping and rezipping cycles of the pre-assembled SNARE complexes^30^. They also measured that the SNARE complexes mostly unzipped within a few seconds at 14pN in the absence of Cpx, but the addition of CpxI extended τ_unzip_ (the latency to unzipping) to hundreds of seconds. However, the SNARE complexes are not functional in the interaction of the CpxI and the pre-assembled SNARE. Yet, it is still unclear how Cpx interacts with SNARE complex in a functional state. Here, we study the interplay of CpxI with individual functional SNARE complex during dynamic transition under constant average force at a fixed trap separation through dual-trap optical tweezers^31, 32^.

Single molecule experiment provides direct evidence on CpxI’s function, and the distinct roles in membrane fusion. In this condition, the SNARE maintained a uniformly distributed four-state transition and the CpxI binding site can be identified. However, it is still illusive how the CpxI binds to the single molecular SNARE complex and interplays with the complete zippering of SNARE. Will the four-helix bundle of SNARE complex be stabilized by the CpxI? Here, we apply the single molecule experiment with dual-trap optical tweezers to understand those mysterious processes involved in the SNARE complex assembly chaperoned by the CpxI molecule.

## Results

### 1. CpxI 1-83aa stabilizes the SNARE complex linker–open state

To observe reversible and regulatory SNARE assembly, we designed SNARE complexes containing the full cytoplasmic domain and a crosslinking site between syntaxin and VAMP2(synaptobrevin-2) near the −6 hydrophobic layer. The SNARE proteins were purified independently, then assembled into SNARE complexes in vitro. The 2,260-bp DNA handle containing an activated thiol group at its 5’end was added to the solution of SNARE complex, at a molar ratio of 1:20. Intramolecular and intermolecular crosslinking occurred in open air between the cysteine residues on syntaxin and VAMP2 and between VAMP2 and the DNA handle. The DNA handle also contains two digoxigenin moieties at 3’ end. Both the thiol group and digoxigenin moieties on the handle were introduced in the PCR through PCR primers. During the single molecule experiment, the SNARE complex was tethered through DNA handle between the anti-dig beads and streptavidin beads by their dual-dig and biotin tag [Figure 1A].

**Figure 1.**
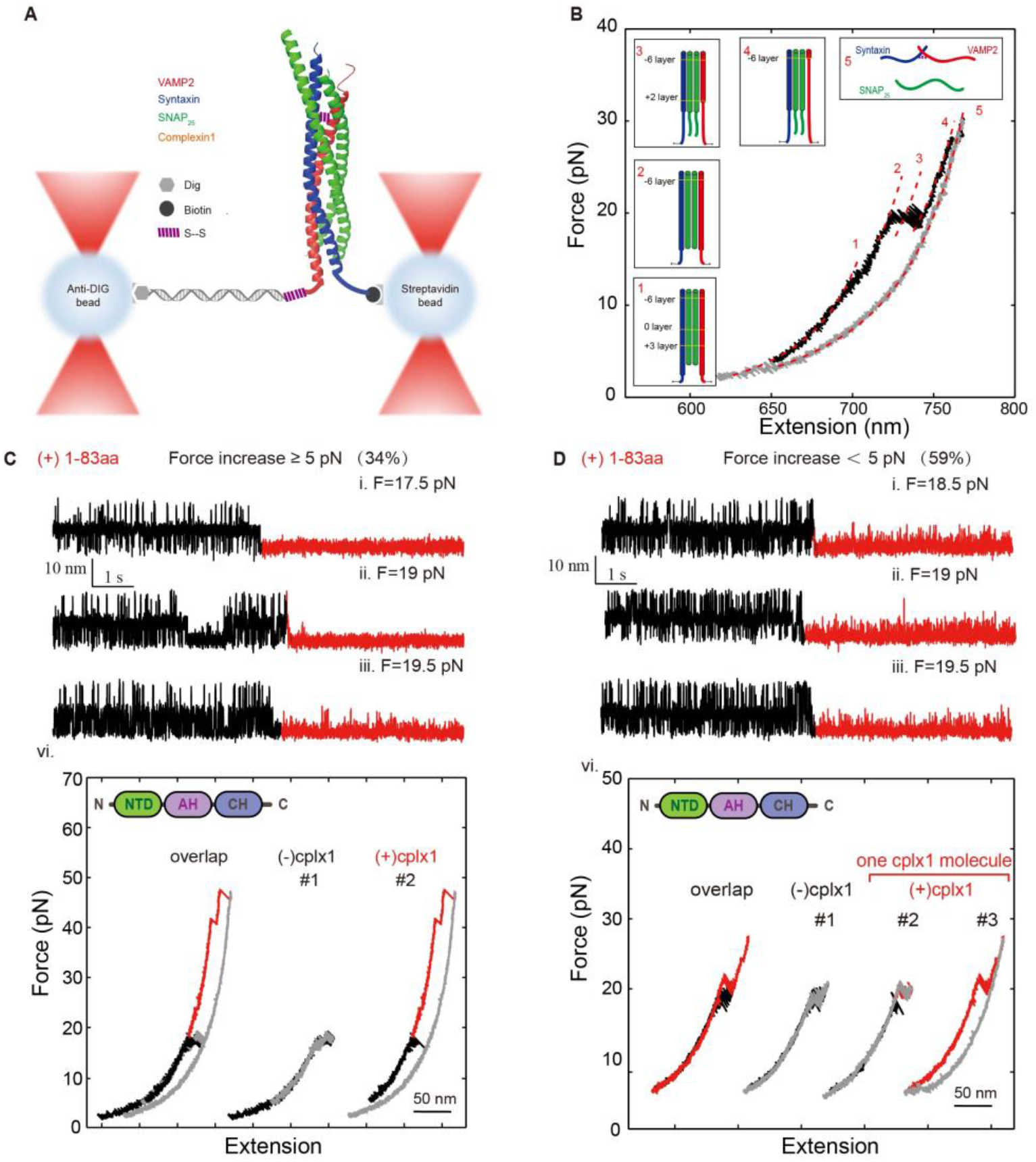
Cpx 1-83aa can stabilize the linker-open state. (A) Schematics of the experimental essay. (B) Force-extension curves (FECs) of a single SNARE complex obtained by pulling (black) and then relaxing (gray) the complex. The continuous regions of the FECs corresponding to different assembly states (illustrated in the insets and marked by red numbers) were fitted by the worm-like chain model (red lines). (CD) Extension-time trajectories (i-iii) and FECs (vi) of single SNARE complexes under constant forces showing SNARE unfolding kinetics before (black) and after (red) the addition of 8 μM 1-83aa in real time. (C, vi) In these liner-open state stabilized signal, 34% of their force-extension curve (FEC) changed dramatically that the unzipped force of C-terminal of SNARE increased by more than 5pN, (D, vi) 59% of their FEC changed slightly that the unzipped force of C-terminal of SNARE increased up to 5pN.

We use dual-trap optical tweezers to capture two beads. One trap is kept stationary, while the other is movable to form a tether of the DNA handle connected with the protein under study. To manipulate a single SNARE complex, we either pulled or relaxed the complex by moving one optical trap relative to the other at a speed of 20 nm/s or held the complex under constant average force at a fixed trap separation. Both force and extension of the protein-DNA tether were recorded at 10 kHz. Specifically, for a reversible two-state transition, the folding energy of the associated protein domain can be measured based on the mechanical work required to unfold the domain, which is equal to the equilibrium force multiplied by the extension change associated with the transition. When gradually pulled, a tethered SNARE complex goes through stepwise zippering of slow association between linker domain, C-terminal domains (CTD), N-terminal domains (NTD) of t- and v-SNAREs, and irreversible dissociation of SNAP25 (synaptosome-associated protein 25) [Figure 1B]. The time-dependent extension and force were mean-filtered using a time window of 7 ms. The Force-extension curve of the handle DNA would be fitted using Worm-like chain model.

To characterize CpxI-dependent SNARE assembly/disassembly, we measured the extension-time trajectories at a mean force where SNARE transits among 3 states, and studied whether the introduction of Cpx or its truncations would interfere with the SNARE transition dynamics. In our tweezer’s configuration, we supplied 8 μM of Cpx in a separate channel (protein channel’) and directly injected to the region where the SNARE folding/unfolding took place. Remarkably, we found that the dynamics extension transition signals of SNARE folding in presence of 1-83aa were the most consistent among all the kind of CpxI. In the presence of 8μM CpxI 1-83aa 79% of 62 transition-state SNARE complex molecules were stabilized at linker-open state or clamped into only N-terminal transition after the addition of CpxI. Interestingly, 94% of their extension-time traces were stabilized at linker-open state, and higher force was required to break the linker-open state again. In these liner-open state stabilized signals, 34% of their force-extension curves (FEC) changed dramatically in which the unzipping force of C-terminal of SNARE increased more than 5pN [Figure 1C], and 59% of their FEC changed slightly that the unzipping force of C-terminal of SNARE increased up to 5pN [Figure 1D]. The 1-83aa domain of CpxI could stabilize the linker-open state of SNARE. Moreover, a different disassembly event appears in consecutive rounds when pulling and relaxing the same SNARE complex, and the probability to detect a force-increased C-terminal disassembly event after another was over 80%, suggesting a strong correlation between the different disassemble events mediated by CpxI. An appealing interpretation for the correlation is that the same CpxI molecular mediated the consecutive disassembly events without dissociation from the SNARE complex during its multiple rounds of reassembly and disassembly.

### 2. NTD in Cpx stabilizes the four-helix bundle of SNARE complex

Previous studies showed that Cpx requires the NTD (1-26 aa) to stabilize the C-terminal of SNARE complex (Choi et al., 2016; Lohmann and Paris, 1968; Xue et al., 2010). Especially, Cpx activates Ca^2+^-triggered neurotransmitter release. Interestingly, CpxI’s 1-83aa may localize to the point where trans-SNARE complexes insert into the fusing membranes^33^, or can position the C-terminal of SNARE complex and lock this domain in the absence of Ca^2+^ and the Ca^2+^ sensor Synaptotagmin 1^34, 35^. These findings suggest that the ability of CpxI to stabilize SNARE complex may mostly depend on its NTD. To pinpoint the possible role of the NTD in the CpxI-dependent SNARE disassembly, we removed the NTD in the CpxI construct and repeated the above experiments. The NTD removal led to significant changes in SNARE complex disassembly. In the presence of 8μM CpxI, 55% of 66 transition-state molecule changed their state after the addition of Cpx. Interestingly, the extension-time traces of 66% of these molecules were still stabilized at linker-open state [Figure 2A], while 18% of their C-terminal transition were blocked [Figure 2B]. As for the latter signal, most SNARE complex couldn’t achieve complete zippering even the force dropped below 5 pN. The rate of linker-open state decreased dramatically, showing that the NTD removal caused significant changes in the SNARE disassembly [Figure 2C], which indicates that the ability of CpxI to stabilize the SNARE complex critically depends on its N-terminal domain.

**Figure 2.**
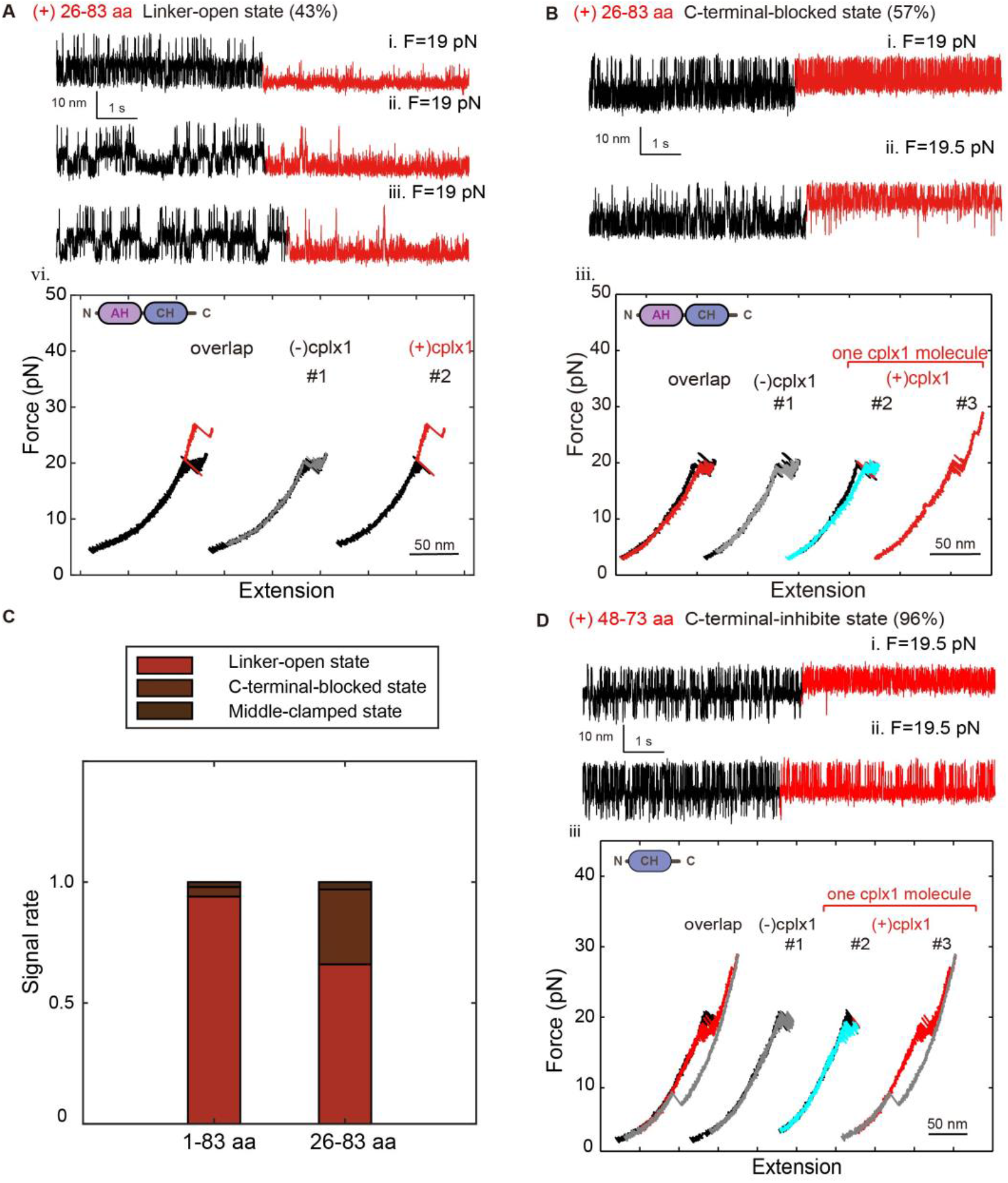
The stabilize function of Cpx NTD and Cpx CH can slightly inhibit the assembly of C-terminal of SNARE complex. In the presence of 8μM, 65.5% of 32 transition-state molecule changed their state after the addition of Cpx. (A) Extension-time trajectories (i-iii) and FECs (vi) of single SNARE complexes under constant forces showing that 43% of them was stabilized to linker-open state after (red) the addition of 8 μM 26-83aa in real time. (B) Extension-time trajectories (i-ii) and FECs (iii) of single SNARE complexes under constant forces showing that 57% of them was locked to the C-terminal blocked state after (red) the addition of 8 μM 26-83aa in real time; relaxing FEC after the addition (cyan). (C) Summary date of the distribution of the different kind of signals introduced by 1-83aa or 26-83aa of Cpx. (D) 81% of 32 transition-state molecule changed their state after the addition of 8 μM 48-73aa of Cpx. Extension-time trajectories (i-ii) and FECs (iii) of single SNARE complexes under constant forces showing that 96% of them was locked to the C-terminal blocked state after (red) the addition; relaxing FEC after the addition (cyan).

The CH domain of Cpx is believed to be the smallest fragment that directly binds with SNARE complex^33^. We supplied 8μM 48-73aa (smallest CH domain) of CpxI to the single-molecule SNARE complex experiment, and found that the CpxI CH slightly inhibits the assembly of C-terminal of SNARE complex. Specifically, with a total of 32 transition-state molecules, 80 % changed their state after the addition of Cpx, and 96% of their C-terminal transition were slightly blocked [Figure 2D]. However, it is noteworthy that these molecules showed full assembly as forces dropped (the cyan line in Figure 2D iii), were quite different from the effect of other longer CpxI fragments.

### 3. CpxI CTD inhibits the assembly of C-terminal and stabilizes the N-terminal of SNARE complex

Recent structure function analyses with C. elegans Cpx revealed that membrane binding is important but not sufficient for Cpx inhibitory effects^28, 29^, leaving room for unknown interactions of the Cpx C-terminus that are instrumental for arresting vesicle fusion. We introduced CTD to figure out whether CTD is functional in the absence of phospholipid. CTD can’t directly bind to SNARE for the lack of CH, so we introduce CTD fragment (83-134aa) with 8μM 1-83aa. Unexpectedly, 56% of 32 transition-state molecules changed their states after the addition of 8μM 1-83aa and 8μM 83-134aa (with molar ratio of 1:1); 62% of 34 transition-state molecules changed their state after the addition of 8μM 1-83aa and 16μM 83-134aa (molar ratio 1:2); 67% of 33 transition-state molecules changed their state after the addition of 8μM 1-83aa and 24μM 83-134aa (molar ratio 1:3) [Figure 3A-C]. Remarkably, more SNARE complexes were clamped to a new state: they can’t reassemble to a fully assembled complex, neither need higher force to break the half-zippered state. Interestingly, the proportion of C-terminal-blocked state and middle-clamped state increases with the concentration of CTD, meanwhile the proportion of the linker-open state decreases [Figure 3D]. Furthermore, the clamping function of Cpx can be reconstituted by 1-83aa of Cpx and its CTD as separate fragments in vitro [Figure 3D, last two columns]. Thus, physical continuity through the length of Cpx is not required and composite Cpx domains can act in tandem to establish a fully functional ensemble. CTD of Cpx seemed to inhibit the full zipper of SNARE complex, but stabilize the half-zipper state. An attractive explanation can be derived from vesicular release experiment of mouse chromaffin cells, the Cpx CTD exhibits a high degree of structural similarity to the C-terminal half of the SNAP25-SN1 domain and lowers the rate of SNARE complex formation in vitro.

**Figure 3.**
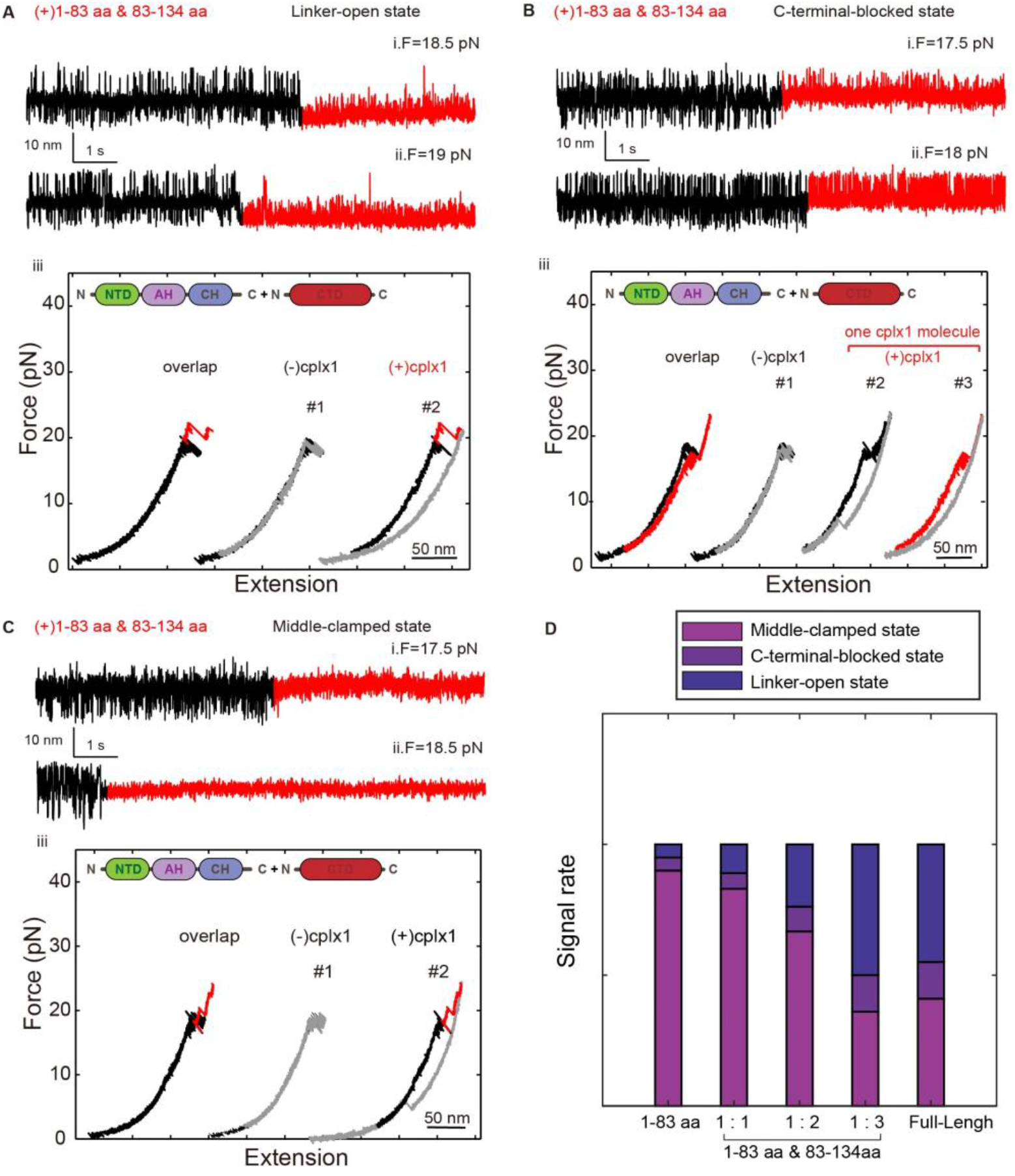
Cpx CTD can inhibit the assemble of C-terminal of SNARE complex and stabilize the N-terminal of SNARE complex. (A-C) Extension-time trajectories (i-ii) and FECs (iii) of single SNARE complexes under constant forces showing SNARE unfolding kinetics before (black) and after (red) the addition of 8 μM 1-83aa and 8/16/24 μM 83-134aa in real time. (D) Summary date of the distribution of the different kind of signals introduced by 1-83aa of Cpx, full-length Cpx, and mixture of 1-83aa and 83-134aa of Cpx. With the increase of CTD’s concentration, the proportion of C-terminal-blocked state and middle-clamped state increase, and meanwhile the proportion of the linker-open state decrease.

Moreover, Cpx: SNAP25-SN1 (C-terminal half) chimeras fully restore function in Cpx deficient cells. Collectively, these results provide evidence to establish a new model wherein the Cpx C-terminus competes with SNAP25-SN1 for binding to the SNARE complex and thereby halts progressive SNARE complex formation before the triggering of Ca^2+^-stimulus^36^. The intramolecular distance between the central domain of CpxI and the far CTD appears to be long enough for the latter to fold back on membrane proximal parts of the SNARE complex and thus may hinder its further zippering. This notion is further supported by the observation that Cpx Cys105 positioned near the central ionic layer of the SNARE complex, leaving the downstream CTD region free to interact with the C-terminal layers of SNARE complex^37^. As a result, the CTD of CpxI hinders SNARE complex assembly.

Since CpxII binds to binary as well as ternary complexes with high affinity^38^, it remains to be shown at which particular step the protein interferes with SNARE assembly. These in vitro results agree with previous reports, showing that full-length Cpx inhibits SNARE-mediated liposome fusion, whereas a truncated variant (amino acid 26-83) does not^6, 39^. Given that CpxII CTD interferes with SNARE zippering, one might expect a retardation of synchronous exocytosis. Similarly, one might ask why the rescue of the function of full-length Cpx need a high concentration addition of Cpx CTD. An attractive explanation from single molecule FRET measurements, suggests that the Cys105 within the Cpx CTD produces only broad FRET efficiency peaks with the labeling site on the SNARE complex (Syntaxin 228,^37^). This indicates conformational variability or motional averaging and suggests that even in the presence of endogenous Cpx the interaction of the CTD with SNAREs is transient and local concentrations of the CTD may not be saturating. Therefore, addition of the CTD peptide is able to enhance the inhibitory function of Cpx.

### 4. Three signals of the FL Cpx-regulated assembly / disassembly of SNARE

Then we introduced full-length of CpxI to interplay with individual SNARE complex held by dual-trap optical tweezers under constant average force with a fixed trap separation (Ma et al., 2015). With the addition of 8μM full-length CpxI, 54% of 74 transition-state molecules changed their state after the addition of CpxI. For 41% of them, the assemble and disassemble process of SNARE complex were limited to the N-terminal (N-terminal transition), suggesting that CpxI might insert into the C-terminal of SNARE Complex to inhibit the complete zippering of SNARE [Figure 4A]. There is also a type of signal (14%) showing that SNARE complex was clamped into the exactly half-zippered state [Figure 4B]. Another kind of data also occupied a considerable part in our data [Figure 4C]. This kind of data showed that SNARE complex maintained in linker-open state after incubation with CpxI, which suggested that CpxI might have the ability to stabilize the four-helix SNARE complex. This ability is necessary for CpxI to help SNARE complex zipper when the Ca^2+^ arrived in a physiological state.

**Figure 4.**
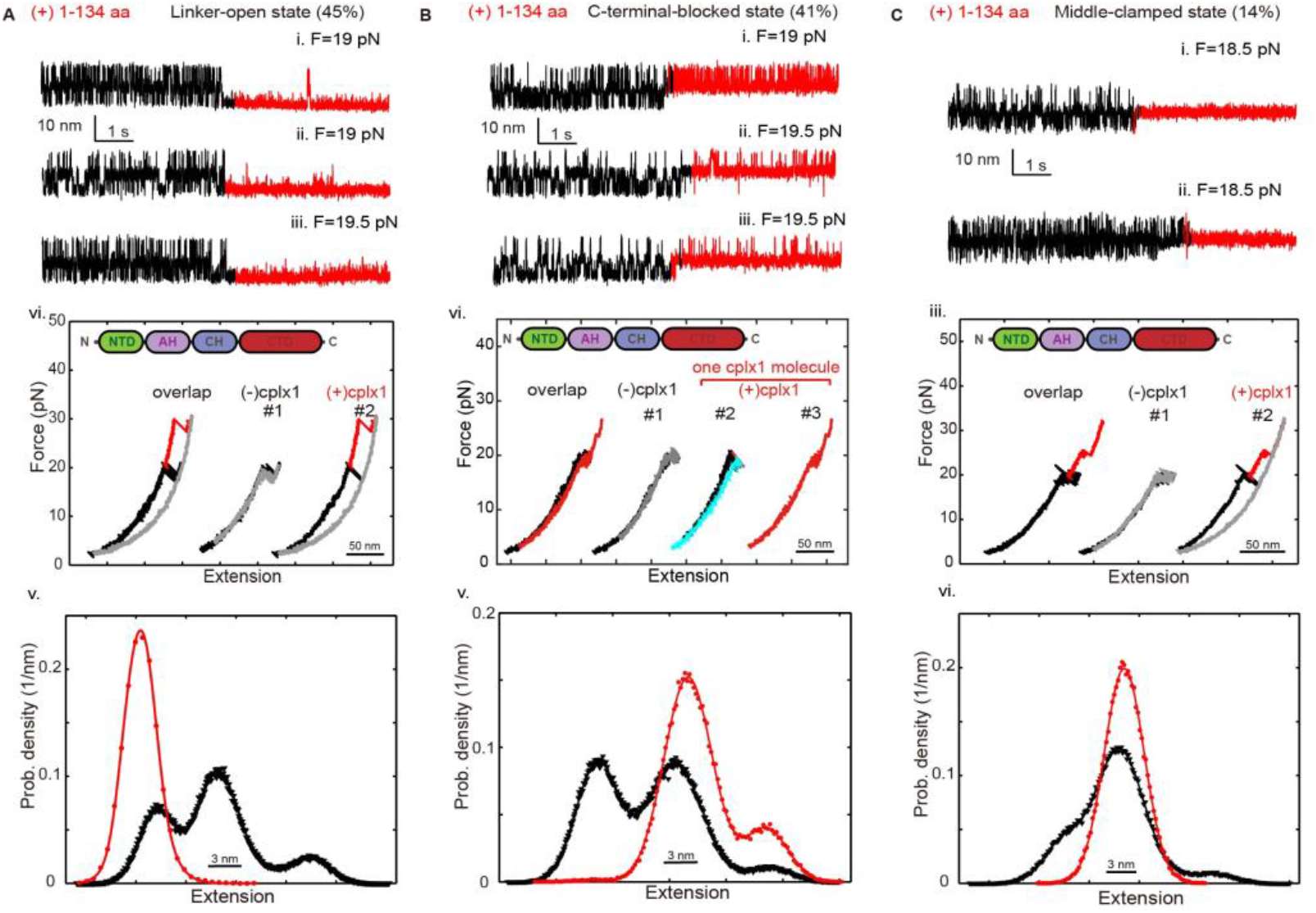
Three signals of the FL Cpx-regulated assembly / disassembly of SNARE. (A) Extension-time trajectories (i-iii), FECs (vi) and HMM fitting (v) of single SNARE complexes under constant forces showing that 45% of them was stabilized to linker-open state after (red) the addition of 8 μM 1-134aa in real time. (B) Extension-time trajectories (i-iii), FECs (vi) and HMM fitting (v) of single SNARE complexes under constant forces showing that 41% of them was locked to the C-terminal blocked state after (red) the addition of 8 μM 1-134aa in real time. (C) Extension-time trajectories (i-ii), FECs (iii) and HMM fitting (vi) of single SNARE complexes under constant forces showing that 14% of them was locked to the middle-clamped state after (red) the addition of 8 μM 1-134aa in real time.

These variable signals were due to synergistic and antagonism of multiple domains. The multiple parts of CpxI work together on a single SNARE complex to assure CpxI a molecular switch – clamp the SNARE complex into half-zippered state without Ca^2+^ to arrest vesicles in the docked state allowing for an appropriate stimulus-secretion coupling, and then help SNARE complex complete assembly in response to a triggering Ca^2+^-stimulus.

## Discussion

The Ca^2+^-triggered exocytosis of neurotransmitters and hormones is a tightly controlled process that has evolved to meet temporal precision and speed of intercellular communication. The core membrane fusion machinery is constituted by a set of three highly conserved proteins known as the SNAREs. Cpx is likely the most controversially discussed SNARE-interacting proteins involved in exocytosis. By applying high-resolution optical tweezers, we had measured the significant different folding energies and kinetics of SNARE complexes in the absence and presence of both wild-type and mutant/truncated CpxI. Here, we showed that CpxI, especially the 1-83aa, can stabilize the four-helix bundle of SNARE motifs. In addition, we found that crucial inhibitory actions of CpxI’s CTD are not only due to membrane binding or vesicle association, but are mediated by direct interaction with SNARE complex. In close correlation, we showed that separate 1-83aa and C-terminal domains of CpxI can efficiently reconstitute the inhibitory signal of the full-length CpxI. Collectively, our results delivered new insight into fundamental mechanisms that constitute the molecular clamp of Ca^2+^-triggered exocytosis.

We found that CpxI’s multiple domains have different effect on the dynamic zippering of SNARE complexes. To examine effect of CpxI on SNARE zippering, we observed a series of prominent long-dwelling states in the middle of fast SNARE transition before and after the introduced of 8 μM full-length and different kind of truncated CpxI. First, we found that the 1-83aa of CpxI can powerfully stabilize the four helix-bundle of SNARE complex. At the same time, no effect on the linker domain of SNARE complex. Second, we confirmed that the ability of CpxI to stabilize the SNARE complex critically depended on its N-terminal domain. Third, we observed a weak and may wrong interaction occur between only CH and the C-terminal of SNARE complex, for the proper and functional orientation need the cooperation of other domains of CpxI. Fourth, we showed that that separate 1-83aa and C-terminal domains of CpxI can efficiently reconstitute the inhibitory signal of the full-length CpxI. Finally, multiple domains work together to make sure a full-length Cpx work as a molecule switch — stabilize the four-helix SNARE complex without Ca^2+^; help SNARE complex zipper when the Ca^2+^ arrived in a physiological state. For the precise realize of this function in vivo, a couple of other regulators (e.g., synaptotagmin) and phospholipid may participate in this process, which need further research.

Experimental results from other workers previously shown that the CTD of CpxII exerts a fusion clamping function, although the underlying mechanisms remained largely enigmatic^25–27, 40^. Although, experiments at the NMJ of C. elegans and murine cortical neurons suggested that the CTD of Cpx guides the protein to vesicular membranes in a curvature sensitive fashion and thus concentrates other inhibitory domains of CpxII at the site of exocytosis for fusion clamping^21, 28^. Some studies counter the hypothesis that the CTD simply targets CpxII to the membrane, but rather suggest a direct blockade of SNARE zippering. In our study, 1-83aa of Cpx was short of the ability the block the C-terminal zippering of SNARE complex compared with full-length Cpx, for lacking CTD. In the same line, our complementation experiments illustrate that separate N-and C-terminal domains of Cpx can efficiently reconstitute the inhibitory signal of the full-length Cpx. For there is no phospholipid membrane or Ca^2+^ in our experiment environment, these results contrast the idea that vesicular targeting of CpxII by its CTD is essential for fusion clamping in chromaffin cells and indicate that the far CTD acts as an independent inhibitory module within the fusion machinery. Collectively, these results are difficult to reconcile with a vesicle targeting role of the CTD of CpxI and rather suggest that this protein region plays an active role in fusion inhibition. Clearly, our observations do not exclude the possibility that vesicular localization is still functionally relevant as it concentrates CpxI at the sites of vesicle fusion.

## Conclusion

Our single molecule optical tweezers experiments have pinpointed that the 26-83aa (AH-CH) of CpxI is the ‘minimal clamping domain’ of Cpx^41^, and Cpx NTD have critical impact in the stable function of Cpx, when Cpx CTD plays an active role in fusion inhibition. In contrast to the stretching of the pre-assembled SNARE complex chaperoned with CpxI in magnetic tweezers experiment^30^, single molecule optical tweezes experiment allows the study of the folding dynamics of SNARE complex in a functional state. Additionally, the dual-trap optical tweezers allow recording of more functional states spanning in the range of 5-40 pN. In vitro analyses in Hela cells by Rothman and colleagues demarcated a region comprising amino acids 26–83 of CpxI as the ‘minimal clamping domain’ of the protein^41, 42^, which is further corroborated by our single molecule studies. Finally, our study suggests that the C-terminal domain (CTD) of CpxI plays an active role in fusion inhibition, also is consistent with recent studies^36^. With the addition of CpxI, assembly and disassembly of SNARE complex is limited to the N-terminal, suggesting that the CpxI binds on the C-terminal of SNARE complex to inhibit the complete zippering of SNARE. Another kind of SNARE complex is switched to the linker-open state, suggesting that the CpxI can stabilize the four-helix bundle of SNARE complex. Therefore, full-length CpxI performs as a molecular switch – stabilizes the four-helix SNARE complex without Ca^2+^, and helps SNARE complex zipper when the Ca^2+^ arrived in a physiological state.

## Materials and method

### Protein purification, labeling

The synaptic SNARE complex consists of VAMP2 (1-92, C2A, Q36C), syntaxin 1 (172-265, C173A, L209C), and SNAP25 (1-206). Substitution or truncation mutations in SNARE proteins and Cpx (C105A, 1-83, 1-73, 26-83, 26-134, 83-134, 73-134) were generated by overlap extension polymerase chain reaction (PCR) using respective primers containing the desired non-homologous sequences^43^. All mutations were confirmed by DNA sequence analysis (BioSune, China). Genes corresponding to syntaxin and VAMP2 and the above Cpx were inserted into pET-SUMO vectors through TA-cloning. The proteins were then expressed in E. coli BL21(DE3) cells and purified as described in the manual of Champion™ pET SUMO Expression System (Invitrogen). Typically, E. coli cell pellets were resuspended in 25mM HEPES, 400 mM KCl, 10% Glycerol, 10mM imidazole, pH 7.7 (25mM HEPES, 800mM NaCl, 5% Glycerol, 2mM β-Me, 0.2% Triton X-100, pH 7.5 for Cpx) and broken up by sonication on ice to obtain clear cell lysates. The lysates were then cleared by centrifugation. The SNARE proteins in the supernatant were bound to Ni-NTA resin and washed by increasing imidazole concentrations up to 20mM. The syntaxin protein was biotinylated in vitro by biotin ligase enzyme (BirA) as described (Avidity, CO). Finally, the His-SUMO tags on both proteins were cleaved directly on Ni-NTA resin by incubating the tagged SNARE proteins/resin slurry with SUMO protease (with a protein-to-protease mass ratio of 100:1) at 4 °C overnight. The SNARE proteins were collected in the flow-through while the His-SUMO tag was retained on the resin. SNAP25 was expressed from the pET-28a vector and purified through its N-terminal His-tag. All SNARE proteins were purified in the presence of 2 mM TCEP to avoid unwanted crosslinking. Peptides of Cpx (48-73,48-83) were synthetized by Yochem Biotech.

### SNARE complex formation and crosslinking

Ternary SNARE complexes were formed by mixing syntaxin, SNAP25 and VAMP2 proteins with 3:4:5 molar ratios in 25 mM HEPES, 150 mM NaCl, 2mM TCEP, pH 7.7 and incubating the mixture at 4 °C for 30mins. Formation of the ternary complex was confirmed by SDS polyacrylamide gel electrophoresis. Excessive SNARE monomers or binary complexes were removed from the ternary complex by further purification through Ni-NTA resin using the His-Tag on the SNAP25 molecule. Middle intramolecular crosslinking around the −6-layer occurred at 34°C, 300rpm, 16 h at a low concentration in 100 mM phosphate buffer, 0.5 M NaCl, pH 8.5 without TCEP. Then the 2,260-bp DNA handle containing an activated thiol group at its 5’ end was added to the solution of SNARE complex, which was just concentrated to more than 70 μM, with a SNARE complex to DNA handle molar ratio of 20:1. Intermolecular crosslinking occurred in open air between VAMP2 and the DNA handle, respectively, in 100 mM phosphate buffer, 0.5 M NaCl, pH 8.5. The DNA handle also contains two digoxigenin moieties at the other end. Both the thiol group and Digoxigenin moieties on the handle were introduced in the PCR reaction through PCR primers. The excess of SNARE complexes in the crosslinking mixture was removed after the SNARE complex-DNA conjugates were bound to the anti-digoxigenin coated beads.

### High-resolution optical tweezers experiments

The high-resolution dual-trap optical tweezers were built by splitting a 1064 nm infrared laser (Spectra Physics) with orthogonal polarization. The optical tweezers were installed on the basement of the Molecular Imaging System in National Facility for Protein Sciences Shanghai with concrete background to isolate from the low-frequency vibration. The instrument was isolated from all the environmental noise including the fan from the laser controller, and all the operation were performed outside the isolated room after the samples were loaded to the microfluidics. Specifically, one trap was kept fixed, while the other is steered by a piezo mirror (MadCity labs). Experiments were carried out at room temperature (22 °C) in the HEPES buffer (25 mM of HEPES, 50 mM NaCl, 0.02% CA630, pH 7.4), supplemented with oxygen scavenging system^32, 44^ Two types of beads are treated separately. The first anti-digoxigenin antibody-coated polystyrene bead (diameter 2.12 um) suspension was mixed with an aliquot of the mixture (crosslinked DNA handle and the SNARE complex), and the first bead is held in the right optical trap; the second bead (diameter 1.76 um) coated with Streptavidin, was subsequently captured in another optical trap and brought close to the first bead to form the protein-DNA tether between two beads. Data were acquired at 20 kHz, mean-filtered online to 10 kHz, and saved on a hard disk for further analysis. In the pulling-relaxation experiment, single SNARE complexes were pulled (relaxed) by increasing (decreasing) the separation between the two optical traps at a constant speed of 10 nm/sec.

#### FEC Analysis of Contour length

To characterize the change in contour length of the SNARE-Cpx complex, unfolding and refolding traces (FECs) were fitted to the wormlike chain model. For the DNA handle, the extension *x_DNA_* as a function of the stretching force *F* is described by the Marko-Siggia formula^45, 46^

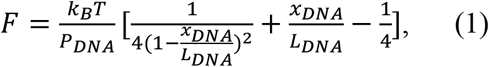

where *P_DNA_* is the persistence length of DNA, *L_DNA_* is the contour length of DNA and *k_B_* is the Boltzmann constant. Similarly, the function for the unfolded polypeptide portion of SNARE complex can be written as

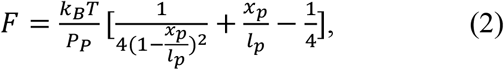

where *F* and *x_P_* are the tension and extension of unfolded polypeptide respectively. The contour length of unfolded polypeptide *l_p_* is related to the number *N* of amino acid in the polypeptide, which is described as *l_p_* =0.4 x *N*, where 0.4 nm is assumed as crystallographic contour length per amino acid^47, 48^. And *P_p_* is the persistence length of unfolded polypeptide. We adopted the constant *k_B_T* = 4.1 *pN · nm* at room temperature.

Therefore, the extension of the protein-DNA handle tether, *X,* is the sum of the extensions of the DNA handle, *x_DNA_*, the unfolded polypeptide portion of SNARE complex, *x_p_*, and the core structure, *h*, i.e.,

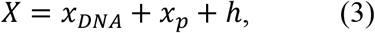

Where *h* is assumed as the spatial length of the folded portion (such as coiled coil) projected along pulling direction, which also makes a contribution to the final extension. In the case of fully folded state (native state) *h*_0_ = 2 nm is determined from the x-ray structure of the protein^16^, whereas for the fully unfolded state, h = 0. And when pulled in an axial direction, *h* = 0.15 × *N_s_ nm*, where *N_s_* is the number of amino acids in the folded portion and 0.15 *nm* is the helical rise of an amino acid along the helix axis. According to the experiment, the stretching force *F* and protein-DNA handle tether *X* between the inner faces of two polystyrene beads follow the equation (4).

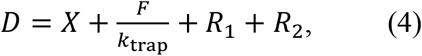

where *k_trap_* = *k*_1_*k*_2_/(*k*_1_ + *k*_2_) is the effective trap stiffness, *R*_1_ and *R*_2_ are the radii of two trapped polystyrene beads and *D* is the trap separation.

The stretching force *F*, the extension *X,* the beads radius *R*_1_ and *R*_2_, the trap stiffness *k*_1_ and *k*_2_, the DNA contour length, *L_DNA_*=768 nm, and the trap separation *D* can be obtained from the experiments, while *h* can be determined from theoretical calculation and crystal structure measurement. Therefore, given the parameters of persistence length of DNA and protein, the structure information of proteins can be derived from fitted contour length of unfolded polypeptide according to equations (1)–(4).

#### Force-extension curve (FEC) fitting

To characterize the folding pathway of protein, the extension of the SNARE complex in each state is calculated by the model. For each state, the extension consists of two parts: the extension of unfolded polypeptide *l_i_r* (*i*= 1, 2, 3, or 4), which can be calculated by the Marko-Siggia formula and the extension of the folded portion. As described above, except for *l_P_, P_DNA_* and *P_P_*, all the other parameters in equations (1)–(4) can be obtained experimentally or theoretically. Therefore, we could fit the experimental extension against the force range of each region (or each state) in FEC, to obtain their best-fit values. The fitting is performed after the FECs are mean-filtered using a time window of 11 ms. The persistence length *P_DNA_* and *P_P_* are different more or less for each molecule, so we firstly confirm these two parameters before each FEC fitting of each sample. The FEC region of fully unfolded state is firstly fitted with *P_DNA_* (10-50 nm) and *P_P_* (0.5-0.9 nm) as fitting parameters, while the known contour length of fully unfolded protein is taken as fixed parameter. Then the best-fit *P_DNA_* and *P_P_* values are utilized as fixed parameters in other FEC regions or states, in which the contour length *l_P_* is to be fitted.

#### Hidden-Markov modeling (HMM) analysis

The HMM analysis was normally performed on the whole extension-time trajectory (typically lasted 1–200 s), after the trajectory was mean-filtered to 5 kHz or 1 kHz. We evaluated the number of states by fitting the histogram distribution of the extension with multiple Gaussian functions. Then, the fitting parameters were further optimized in the hidden Markov model with a four-state transition model using gradient descent. The HMM analyses yielded the corresponding best-fit parameters from the experimental traces, such as the equilibrium force, and the extension change.

## Author Contributions

T. H., N. F., and F. G. purified the protein, and collected the single molecule data; T. H., N. F., L. M., and Y. R. analyzed the data; Y. R. and F. G. built the instrument; J. L. and L. M. provided overall guidance and support; T. H. and Y. R. drafted the manuscript, and all authors revised the manuscript.

## Notes

The authors declare no competing financial interests.

## ACKNOWLEDGMENT

We thank the National Center for Protein Science in Shanghai for use of the dual-trap optical tweezers. T. H. acknowledges Dr. Y. Gao for help on the protein purification, SNARE assembly, and single molecule experiment.

